# Dietary butyrate treatment enhances healthy metabolites by longitudinal untargeted metabolomic analysis in amyotrophic lateral sclerosis mice

**DOI:** 10.1101/2022.01.15.476456

**Authors:** Destiny Ogbu, Yongguo Zhang, Katerina Claud, Yinglin Xia, Jun Sun

## Abstract

Microbial metabolites affect the neuron system and muscle cell functions. Amyotrophic Lateral Sclerosis (ALS) is a multifactorial neuromuscular disease. Our previous study has demonstrated elevated intestinal inflammation and dysfunctional microbiome in ALS patients and an ALS mouse model (human-SOD1^G93A^ transgenic mice). However, the metabolites in ALS progression are unknown. Using an unbiased global metabolomic measurement and targeted measurement, we investigated the longitudinal changes of fecal metabolites in the SOD1^G93A^ mice over the course of 13 weeks. We compared the changes of metabolites and inflammatory response in age-matched WT and SOD1^G93A^ mice treated with bacterial product butyrate. We found changes in carbohydrate levels, amino acid metabolism, and formation of gamma-glutamyl amino acids. Shifts in several microbially-contributed catabolites of aromatic amino acids agree with butyrate-induced changes in composition of gut microbiome. Declines in gamma-glutamyl amino acids in feces may stem from differential expression of GGT in response to butyrate administration. Due to signaling nature of amino acid-derived metabolites, these changes indicate changes in inflammation (e.g. histamine) and contribute to differences in systemic levels of neurotransmitters (e.g. GABA, glutamate). Butyrate treatment was able to restore some of the healthy metabolites in ALS mice. Moreover, microglia in the spinal cord were measured by the IBA1 staining. Butyrate treatment significantly suppressed the IBA1 level in the SOD1^G93A^ mice. The serum IL-17 and LPS were significantly reduced in the butyrate treated SOD1^G93A^ mice. We have demonstrated an inter-organ communications link among metabolites, inflammation, and ALS progression, suggesting the potential to use metabolites as ALS hallmarks and for treatment.

**Graphic Abstract:** We compared the changes of metabolites and inflammatory response in age-matched WT and SOD1^G93A^ mice treated with bacterial product butyrate. Butyrate treatment was able to restore some of the healthy metabolites in ALS mice. Due to signaling nature of amino acid-derived metabolites, these changes indicate changes in inflammation and contribute to differences in systemic levels of neurotransmitters (e.g. GABA, glutamate). Moreover, butyrate treatment significantly suppressed the microglia IBA1 level and aggregated SOD1^G93A^ in the SOD1^G93A^ mice. The inflammatory cytokine, e.g serum IL-17, was significantly reduced in the butyrate treated SOD1^G93A^ mice. We have demonstrated an inter-organ communications link among metabolites, inflammation, and ALS progression, suggesting the potential to use metabolites as ALS hallmarks and for treatment.

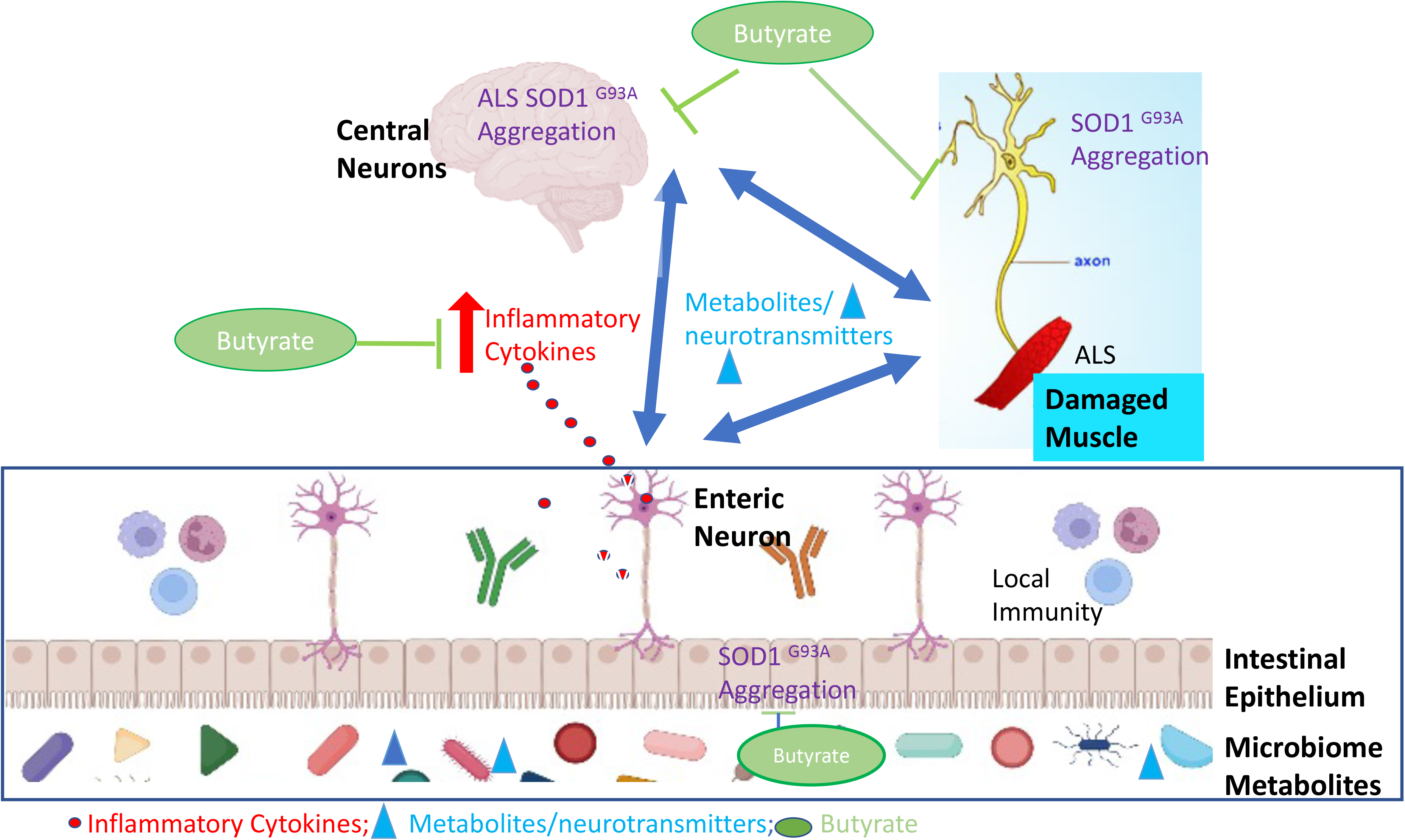

## Introduction

Amyotrophic lateral sclerosis (ALS), or Lou Gehrig’s disease, is a multifactorial neurodegenerative disease characterized by the premature death of motor neurons and has an average survival of three to five years in humans ^1^. Currently, treatment with the FDA approved drug, Riluzole, merely extends the patient’s life span for a few months ^2^. The second drug Radicava (Edavarone) approved in 2017 shows moderate efficacy ^3^. Thus, new treatments are still needed for alleviating disease progression and improving the life quality of patients with ALS.

The inter-organ communications may play important role in the disease progression. Celiac disease with neurologic manifestations was misdiagnosed as ALS ^4–6^. There is a possible link between ALS and sensitivity to gluten ^7^. Moreover, inflammatory cytokines and bacterial product LPS were elevated in ALS ^8, 9^. Autoimmune disease (e.g. Crohn’s disease) associations with ALS indicate the contribution of genetic or environmental risk factors to the pathogenesis and progression of ALS ^10^. Early diagnosis of ALS has been a long-standing challenge in the field. Metabolomics could be potential diagnostic markers for ALS. Human and mouse studies have shown alterations in carbohydrate, amino acid and lipid metabolism within the serum and cerebral spinal fluid ^11–13^. A PNAS study has shown that the altered metabolites in the ALS-like SOD1^G93A^ model is associated with a progressive state of acidosis ^14^. Thus, understanding the metabolomics and intestinal changes will be important for the neuron pathogenesis in ALS.

Studies of metabolism in ALS through the microbiome and microbial metabolites is a growing field. We are the first to report the elevated intestinal inflammation/permeability and reduced beneficial bacteria ^15^. Furthermore, we have shown the beneficial role of bacterial product butyrate in treating dysbiosis, intestinal leakage, and abnormal enteric neurons in the ALS-like SOD1^G93A^ model ^16, 17^. Gut microbe-produced metabolites directly contribute to the ALS disease phenotype ^18^. These findings suggest metabolites may be a potential gut-neuron-microbiome mechanism that drive the ALS pathology. However, studies focusing the metabolites and inflammation in ALS as well as the metabolic changes following butyrate treatment are lacking.

The goal of this study is to characterize metabolic changes in the SOD1^G93A^ model and to gain insights into fundamentals of neuromuscular function and metabolites in the ALS treatment by targeting metabolites. The transgenic mouse model SOD1^G93A^ mice were chosen because a subset of familial cases of ALS are associated with mutation in *SOD1* gene encoding Cu/Zn superoxide dismutase. SOD1^G93A^ model shows neuronal and muscular impairment like human ALS. Moreover, we examined the changes of metabolites from feces of SOD1^G93A^ mice with or without butyrate treatment and correlated with the inflammation and neurological changes. Our data suggest that changes within metabolite profile are correlated with altered inflammatory cytokines and disease progression. Importantly butyrate treatment significantly reduced metabolic differences between the WT and SOD1^G93A^ mice. Better understanding the intestinal dysfunction and metabolites in ALS will help for the early diagnosis and development of new treatments.

## Materials and Methods

### Animals

SOD1^G93A^ and age matched wild type mice were used in this study, SOD1^G93A^ mice were originally purchased from Jackson Laboratory (B6SJL-Tg (SOD1-G93A) 1Gur/J, stock No. 002726). All experiments were carried out in strict accordance with the recommendation in the Guide for the Care and Use of Laboratory Animals of the National Institutes of Health. Mice were provided with water ad libitum and maintained on a 12-hour dark/light cycle. The protocol was approved by the IACUC of University of Illinois at Chicago Committee on Animal Resources (ACC 18-233).

### Butyrate treatment in mice

SOD1^G93A^ mice and age matched wild type mice were divided into two groups randomly, non-treatment group and butyrate treatment group. The butyrate-treated group receives sodium butyrate (Sigma-Aldrich, 303410, St. Louis, MO) at a 2% concentration in filtered drinking water. Control group receives filtered drinking water without sodium butyrate. All animals are weighted and received detailed clinical examination, which included appearance, movement and behavior patterns, skin and hair conditions, eyes and mucous membranes. If restricted outstretching of the hind legs was observed when the tail was held, it means the symptom of ALS is obvious. Lie the mouse on the back, if it can’t turn over in 30 seconds, the mouse is humanely sacrificed.

### Fecal Sample collection

Metabolon received 76 fecal samples from 6 WT and 6 SOD1^G93A^ mice at baseline, 7 weeks, and 14 weeks post butyrate treatment. For untreated animals, fecal samples from 5 WT and 5 SOD1^G93A^ mice were received at 4, 8, 13, and 17 weeks. Global metabolic profiles were determined from the experimental groups.

Bioinformatic Analysis of Metabolomics Data via Metabolon Platform Sample Accessioning and Sample Preparation: **Sample accessioning and** preparation were processed via the Metabolon LIMS system. Following receipt, all fecal samples were inventoried and immediately stored at -80°C until processed. Samples were prepared using the automated MicroLab STAR® system from Hamilton Company.

Before processing extraction for quality control (QC) and quality assurance (QA), several recovery standards were added to remove protein, dissociate small molecules bound to protein or trapped in the precipitated protein matrix, and to recover chemically diverse metabolites, and proteins were precipitated with methanol under vigorous shaking for 2 min (Glen Mills GenoGrinder 2000) followed by centrifugation. The resulting extract was divided into five fractions: two for separate analyses by reverse phase (RP)/UPLC-MS/MS methods with positive ion mode electrospray ionization(ESI)^18^, one for analysis by RP/UPLC-MS/MS with negative ion mode ESI, one for analysis by HILIC/UPLC-MS/MS with negative ion mode ESI, and one sample was reserved for backup. Samples were placed briefly on a TurboVap® (Zymark) to remove the organic solvent. The sample extracts were stored overnight under nitrogen before preparation for analysis.

### QA/QC

First, several types of controls were analyzed in concert with the experimental samples based on the Metabolon QC recovery and Internal standards, including: (1) generated the pooled matrix sample by taking a small volume of each experimental sample to serve as a technical replicate throughout the data set; ^19^ extracted water samples to serve as process blanks; and (3) carefully chose a cocktail of QC standards to ensure it not to interfere with the measurement of endogenous compounds and spiked it into every analyzed sample, allowed instrument performance monitoring and aided chromatographic alignment. Then, instrument variability was determined by calculating the median relative standard deviation (RSD) for the standards that were added to each sample prior to injection into the mass spectrometers. Overall process variability was determined by calculating the median RSD for all endogenous metabolites (i.e., non-instrument standards) present in 100% of the pooled matrix samples. Experimental samples were randomized across the platform run with QC samples spaced evenly among the injections.

### Data Extraction and Compound Identification

Raw metabolomics data were generated by the Ultrahigh Performance Liquid Chromatography-Tandem Mass Spectroscopy (UPLC-MS/MS). All methods utilized a Waters ACQUITY ultra-performance liquid chromatography (UPLC) and a Thermo Scientific Q-Exactive high resolution/accurate mass spectrometer interfaced with a heated electrospray ionization (HESI-II) source and Orbitrap mass analyzer operated at 35,000 mass resolution. The sample extract was dried then reconstituted in solvents compatible to each of the four methods as described in sample preparation. Raw data was extracted, peak-identified and QC processed using Metabolon’s hardware and software. Compounds were identified by comparison to library entries of more than 3300 commercially available purified standard compounds or recurrent unknown entities. Metabolon maintains a library based on authenticated standards that contains the retention time/index (RI), mass to charge ratio (*m/z)*, and chromatographic data (including MS/MS spectral data) on all molecules present in the library. Thus, biochemical identifications are based on three criteria: (1) retention index within a narrow RI window of the proposed identification,^19^ accurate mass match to the library +/- 10 ppm, and (3) the MS/MS forward and reverse scores between the experimental data and authentic standards. The MS/MS scores are based on a comparison of the ions present in the experimental spectrum to the ions present in the library spectrum. While there may be similarities between these molecules based on one of these factors, the use of all three data points can be utilized to distinguish and differentiate biochemicals.

### Curation

To ensure the available data set for statistical analysis and data interpretation have a high quality, a variety of QC and curation procedures were carried out to ensure accurate and consistent identification of true chemical entities, and to remove those representing system artifacts, mis-assignments, and background noise. For example, metabolon data analysis was performed to confirm the consistency of peak identification among the various samples using proprietary visualization and interpretation software. Library matches for each compound were checked for each sample and corrected if necessary.

### Metabolite Quantification and Data Normalization

Peaks were quantified using area-under-the-curve. Data normalization was performed based on whether the studies were required for more than one day of analysis or not. No normalization was performed for those studies that required one day of analysis. For studies spanning multiple days, a data normalization step was performed to correct variation resulting from instrument inter-day tuning differences. Essentially, each compound was corrected in run-day blocks by registering the medians to equal one (1.00) and normalizing each data point proportionately (termed the “block correction”). In certain instances, biochemical data may have been normalized to an additional factor (e.g., cell counts, total protein as determined by Bradford assay, osmolality, etc.) to account for differences in metabolite levels due to differences in the amount of material present in each sample.

### Immunofluorescence

Lumbar spine tissues were freshly isolated and paraffin-embedded after fixation with 10% neutral buffered formalin. Immunofluorescence was performed on paraffin-embedded sections (5 μm). After preparation of the slides as described previously ^20^, tissue samples were incubated with anti-IBA1 antibody (Cell Signaling, 17198, Danvers, MA) at 4°C overnight. Samples were then incubated goat anti-rabbit Alexa Flour 594 (Invitrogen, A32740, Carlsbad, CA) for 1 h at room temperature. Tissues were mounted with Slow Fade (Invitrogen, s2828, Carlsbad, CA), followed by a coverslip, and the edges were sealed to prevent drying. Specimens were examined with Zeiss laser scanning microscope (LSM) 710. Fluorescence intensity was determined by using Image J software. This method determines the corrected total fluorescence by subtracting out background signal, which is useful for comparing the fluorescence intensity between cells or regions.

### Mouse Cytokines

Mouse blood samples were collected by cardiac puncture and placed in tubes containing EDTA (10 mg/mL). Mouse cytokines were measured using a Cytokine & Chemokine Convenience 26-Plex Mouse ProcartaPlex™ Panel 1 (Invitrogen, EPXR260-26088-90, Carlsbad, CA) according to the manufacturer’s instructions. Briefly, beads of defined spectral properties were conjugated to protein-specific capture antibodies and added along with samples (including standards of known protein concentration, control samples, and test samples) into the wells of a filter-bottom microplate, where proteins bound to the capture antibodies over the course of a 2-hour incubation. After washing the beads, protein-specific biotinylated detector antibodies were added and incubated with the beads for 1 hour. After removal of excess biotinylated detector antibodies, the streptavidin-conjugated fluorescent protein R-phycoerythrin was added and allowed to incubate for 30 minutes. After washing to remove unbound streptavidin–R-phycoerythrin, the beads were analyzed with the Luminex detection system (Bio-rad, Bio-Plex 200 Systems, Hercules, CA).

### LPS Detection

LPS in serum samples was measured with limulus amebocyte lysate chromogenic end point assays (Hycult Biotech, HIT302, Plymouth, PA) according to the manufacturer’s indications. The samples were diluted 1:4 with endotoxin-free water and then heated at 75°C for 5 minutes on a warm plate to denature the protein before the reaction. A standard curve was generated and used to calculate the concentrations, which were expressed as EU/mL, in the serum samples.

### Statistical Analysis

Metabolite data were expressed as a fold change ratio of genotype vs. control, all other data are expressed as the means ± SD. All statistical tests were 2- sided. The *p* values < 0.05 were considered statistically significant. For the summary of numbers of biochemicals that achieved approaching statistically significant (0.05<*p*<0.10) were also reported. Standard statistical analyses of metabolite data are performed in Array Studio on log transformed data. For those analyses of metabolite data not standard in Array Studio, the programs R (http://cran.r-project.org/) are used. A Welch’s two-sample t-test was utilized to assess the mean differences of fold-change values between the study groups. To accommodate high within-group variability, a non-parametric Wilcoxon rank sum test was also performed to compare the median differences of fold-change values between the study groups. In this study, Wilcoxon rank sum test returned similar results with Welch’s two-sample t-test. Thus, the biochemical interpretation of this study is based on the results of the Welch’s two-sample t-test. The differences between samples for more than two groups were analyzed using one-way ANOVA or its non-parametric alternative Kruskal-Wallis test based on the data normality or not detected by the Shapiro-Wilks normality test. The *p* values in ANOVA analyses were adjusted for correction of multiple comparisons using the Tukey method to ensure accurate results. Principal component analysis (PCA) was performed to reduce the dimensionality of the data for visually assessing similarities and differences between samples. Random Forest (RF) Analysis was performed to identify the biomarkers differentiating classification groups. RF analysis generated an accompanying list of the top metabolites contributing most to the separation of the groups being compared. All the other statistical analyses except for metabolite data were performed using GraphPad Prism 9.2.0 (GraphPad, Inc., San Diego, CA) and confirmed using the R software (R version 4.0.4, 2021, The R Foundation for Statistical Computing Platform).

## Results

### Altered microbial metabolites in the ALS mice over course of disease

ALS is a progressive disorder, where the onset of muscle weakness spreads to adjacent body regions, ^21^ thus, we evaluated when fecal metabolite profile changes over disease course. We investigated the longitudinal changes of fecal metabolites in ALS progression using SOD1^G93A^ mice at 4-, 8-, 13-, and 17-week-old. There is a total of 797 compounds identified in the samples. We utilized a principal component analysis (PCA) to visually compare the metabolic profile at 4-, 8-, 13- and 17- week-old between untreated WT and SOD1^G93A^ mice (**Fig. 1A**). The PCA did not show sample separation related to time or genotype, which may be attributed to inter-subject variability in masking the effects related to the genotype. However, regardless of the overlapping between datasets or lack of PCA group separation, individual metabolite levels are still significantly different between groups.

**Figure 1.**
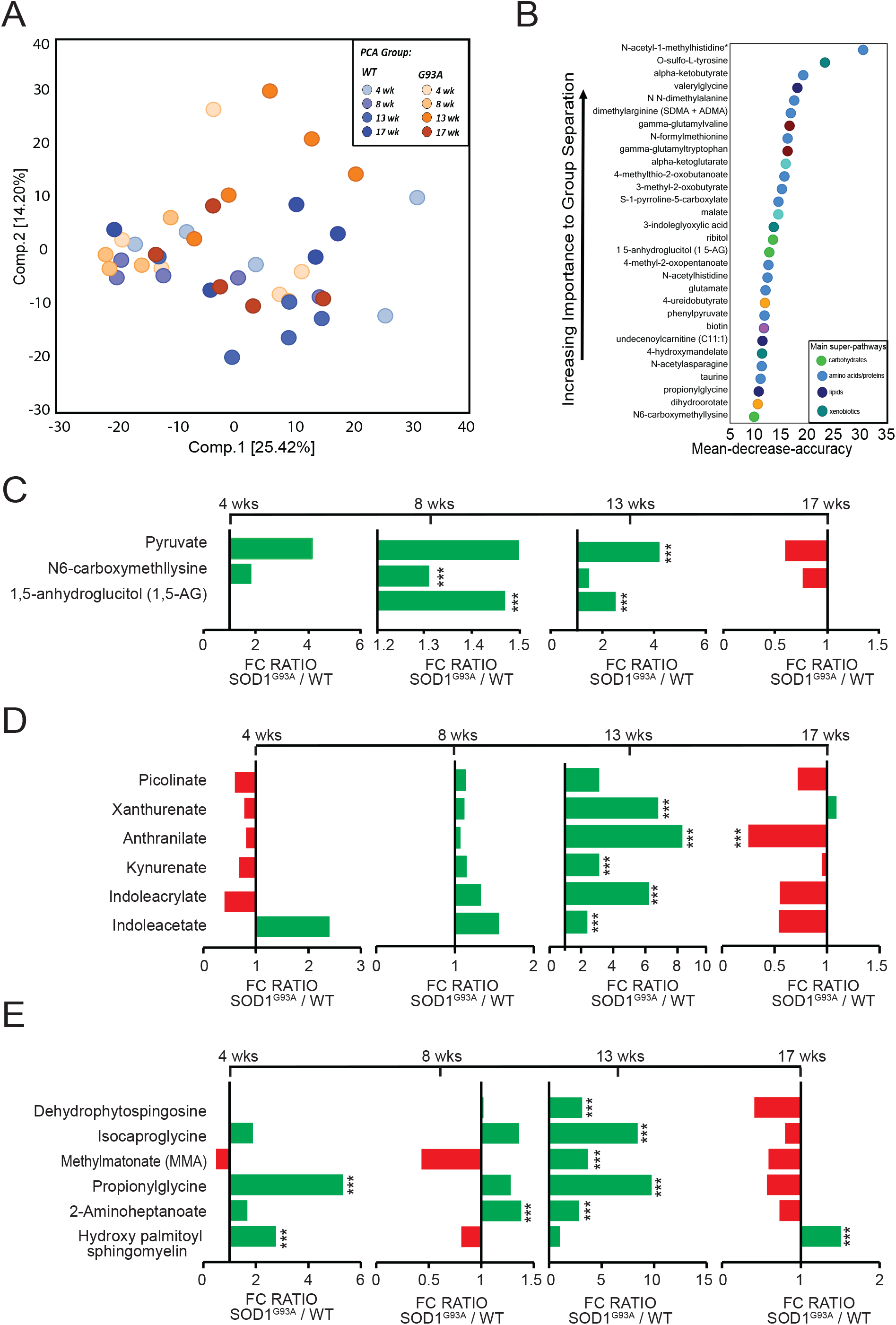
Impact of butyrate treatment on metabolite profile in ALS mice over the course of the disease. (**A**) PCA comparing the biochemical similarity between WT and SOD1^G93A^ mice over the course of disease. (**B**) Random Forest (RF) analysis showing the top 30 most important metabolites that differed between WT and SOD1^G93A^ over the course of disease. The variables are ordered top-to-bottom as most-to-least important. Different color indicates specific metabolites resulting from different super-pathways like green= carbohydrate, light blue= amino acid, blue= lipids, teal= xenobiotics. Fold-change ratio generated by (**C**) carbohydrate (**D**) amino acid (**E**) lipid changes overtime. Differences are assessed by the Welch’s two-sample t-test. WT (N= 6) & SOD1^G93A^ (N=6) mice. *P<0.05, **P<0.01, ***P<0.001

As shown in **Table 1**, SOD1^G93A^ mice experience changes for a total of 258 biochemicals with significance of p ≤0.05 from 4 weeks to 17 weeks of age. We compared the changes in metabolites using a ratio of SOD1^G93A^ to WT mice via Welch’s two sample t-test. At 4 weeks, 61 metabolites were altered (where 49 increased and 12 decreased). At 8 week-old, 7 metabolites changed (of which 5 increased and 2 decreased). SOD1^G93A^ mice had a total of 180 metabolites (of which 175 increased while 5 decreased) change at 13 weeks or onset. Lastly, 10 metabolites changed for SOD1^G93A^ mice at 17 weeks (of which 6 increased and 4 decreased). SOD1^G93A^ mice approached significance with 0.05<p<0.10 for a total of 165 metabolites from 4 weeks to 17 weeks of age. Specifically, at 4-week-old, 54 metabolites were altered in SOD1^G93A^ mice (of which 38 increased while 16 decreased). At 8-week-old, 15 metabolites were altered (8 increased and 7 decreased. 0.05<p<0.10). At 13 weeks, SOD1^G93A^ mice experience the most changes with 72 metabolites altered where 59 increased while 13 decreased. At 17-week-old, 24 metabolites were altered (6 increased and 18 decreased), suggesting the significant changes of metabolites in the progress of the disease in the SOD1^G93A^ mice.

**Table 1.** ALS mice experienced the metabolic alterations over the course of disease compared to WT.

| Welch's Two-Sample t-test | Statistical Comparisons |  |  |  |
| --- | --- | --- | --- | --- |
|  | SOD1<sup>G93A</sup>/ WT |  |  |  |
|  | 4 wk | 8wk | 13 wk | 17 wk |
| Total biochemicals $p \leq 0.05$ | 61 | 7 | 180 | 10 |
| Biochemicals ($\uparrow\downarrow$) | 49 \| 12 | 5 \| 2 | 175 \| 5 | 6 \| 4 |
| Total biochemicals<br>$0.05 < p < 0.10$ | 54 | 15 | 72 | 24 |
| Biochemicals ($\uparrow\downarrow$) | 38 \| 16 | 8 \| 7 | 59 \| 13 | 6 \| 18 |

Next, we observed a random forest (RF) ^22^ analysis of metabolites between WT and SOD1^G93A^ animals which revealed the impact of different metabolites **(Fig. 1B)**. A list of 30 biochemicals contribute to the difference between WT and SOD1^G93A^ mice. The top metabolite groups are amino acids/proteins and xenobiotics. The most significant alterations in metabolites were generated by four super-pathways including (A) carbohydrate, (B) protein/amino acids (C) lipids, and (D) xenobiotics metabolism. The top three metabolites were N-acetyl-1-methylhistidine, O-sulfo-L-tyrosine and alpha ketobutyrate and which were in amino acids and xenobiotics groups, respectively.

#### Changes of carbohydrate in the SOD1^G93A^ mice

Carbohydrate fermentation yields dietary fibers that are a primary source of energy for gut microbiota ^23^. Defects in glucose metabolism may contribute to disease progression in ALS ^24^. **Fig. 1C** shows changes in some glucose-related carbohydrate metabolites for SOD1^G93A^ mice over disease course. SOD1^G93A^ mice had a significant increase in N6- carboxymethyllysine and 1,5-anhydroglucitol (1,5-AG) at 8 weeks. At 13 weeks of age, 1,5-anhydroglucitol and pyruvate significantly increased. Fecal metabolite levels of glucose-related carbohydrates differed between ALS and WT mice.

### Changes of amino acids/proteins in SOD1^G93A^ mice

Tryptophan is an essential amino acid in the microbiome ^25^ and metabolism of tryptophan is altered in ALS pathogenesis ^26^. We observed changes in tryptophan metabolism in ALS mice over disease course (**Fig. 1D**). The tryptophan-related metabolites: xanthurenate, anthranilate, kynurenate, indoleacrylate, and indoleacetate significantly increased at 13 weeks for SOD1^G93A^ mice. Only anthranilate decreased at 17 weeks. suggesting the altered metabolites are associated with ALS onset.

#### Changes of lipids in SOD1^G93A^ mice

Dysfunctions in lipid metabolism, including fatty acids, triacylglycerols, phospholipids, sterol lipids and sphingolipids, have been identified as potential drivers of pathogenesis in ALS ^27^. **Fig. 1E** shows observed changes in sphingomyelins, sphingosines, and fatty acids in SOD1^G93A^ mice. At 4 weeks of age, SOD1^G93A^ mice had an increase in propionylglycine and hydroxy palmitoyl sphingomyelin. 2-aminoheptanoate increased at 8 weeks. At 13 weeks, dehydrophytospingosine, isocaproglycine, methylmatonate ^28^, propionylglycine, and 2-aminoheptanoate significantly increased at 13 weeks of age. Only, hydroxy palmitoyl sphingomyelin increased at 17 weeks of age for SOD1^G93A^ mice. These results depict changes in lipid metabolism during ALS progression.

### Dietary butyrate treatment altered the metabolite profile of ALS mice over disease course

Our previous study has shown that bacterial butyrate delayed ALS progression in the SOD1^G93A^ mice by restoring the intestinal function and microbiome ^16^. Here, we assessed the effect of dietary butyrate on metabolite profile of SOD1^G93A^ and WT mice over the course of 14 weeks. In **Fig. 2A**, we observed a PCA to visually assess similarities between untreated and butyrate-treated groups at baseline, 7 weeks, and 14 weeks. The PCA shows moderate separation between untreated baseline sample compared to butyrate-treated samples at 7 and 14 weeks. This suggests butyrate treatment altered the metabolic profile of WT and SOD1^G93A^ mice.

**Figure 2.**
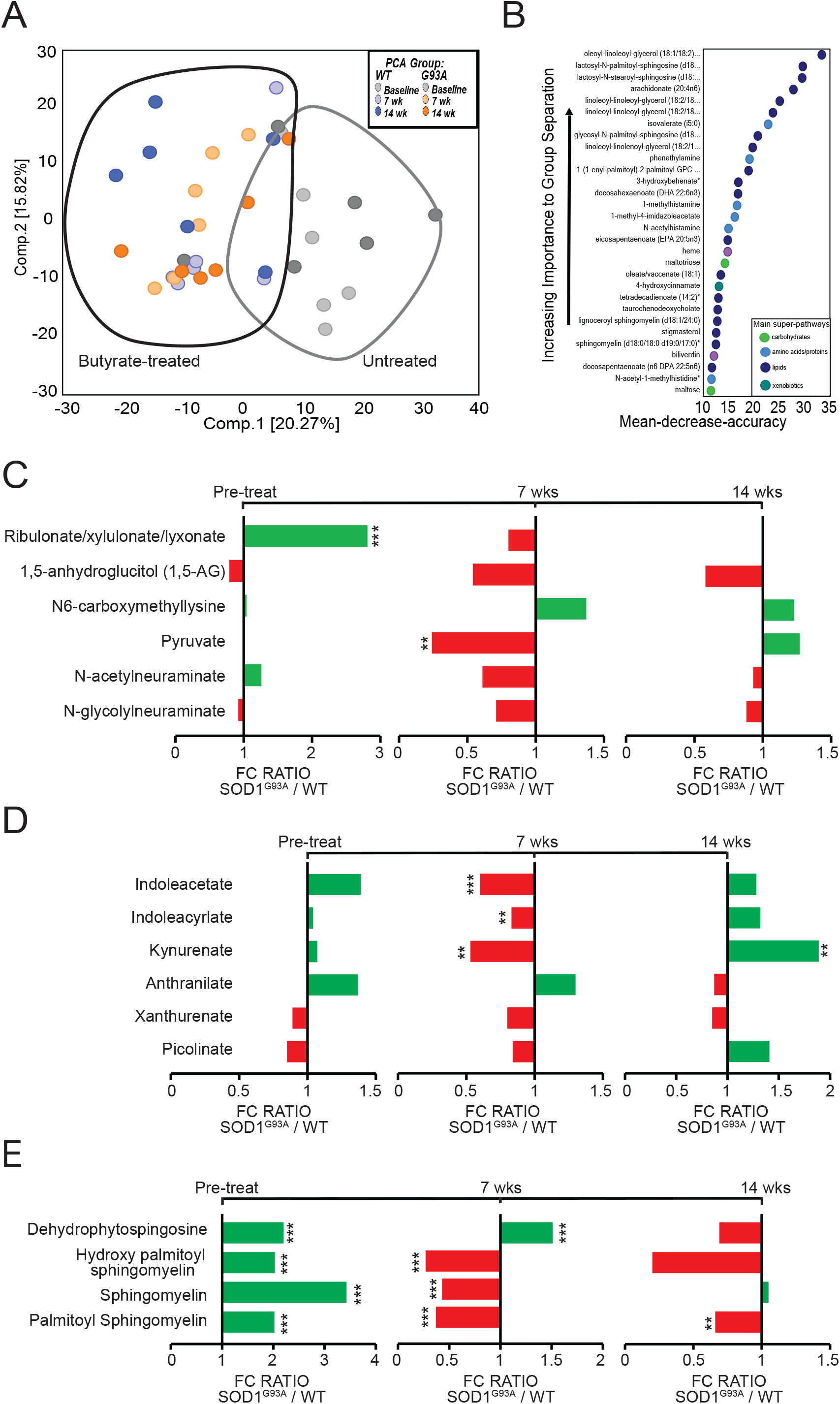
Effect of butyrate treatment on metabolite profile in ALS mice over the course of disease. **(A)** PCA comparing the biochemical similarity between butyrate-treated WT and SOD1^G93A^. **(B)** RF showing the top 30 most important metabolites that differed between longitudinal butyrate treatment WT and SOD1^G93A^ mice. Fold-change ratio generated by **(C)** carbohydrates **(D)** amino acid **(E)** lipid. Differences are assessed by the Welch’s two- sample t-test. WT (N= 6) & SOD1^G93A^ (N=6) mice. *P<0.05, **P<0.01, ***P<0.001

In **Table 2**, SOD1^G93A^ mice achieved statistical significance p ≤ 0.05 for a total of 188 metabolites post-butyrate treatment. At 7-weeks of treatment, 107 metabolites were altered (of which 64 increased and 43 decreased) while 81 metabolites were altered at 14 weeks of treatment (of which 32 increased and 49 decreased). In addition, SOD1^G93A^ animals showed changes for a total of 115 biochemicals with approaching significance of 0.05<p<0.10. At 7- weeks of butyrate treatment, 52 metabolites were altered in SOD1^G93A^ mice (of which 30 increased and 22 decreased). At 14-weeks of butyrate treatment, 63 metabolites were altered (30 increased and 33 decreased) suggesting the significant changes of butyrate treatment in metabolites in the SOD1^G93A^ mice.

**Table 2.** Butyrate (But) treatment contributes to the altered profile of metabolites in ALS mice, compared to WT mice.

| Statistical Comparisons |  |  |  |
| --- | --- | --- | --- |
| Welch's Two-Sample t-Test | SOD1<sup>G93A</sup>/ WT |  |  |
|  | Baseline | 7wk But | 14 wk But |
| Total biochemicals $p \leq 0.05$ | 74 | 107 | 81 |
| Biochemicals ($\uparrow\downarrow$) | 13 \| 61 | 64 \| 43 | 32 \| 49 |
| Total biochemicals $0.05 < p < 0.10$ | 58 | 52 | 63 |
| Biochemicals ($\uparrow\downarrow$) | 20 \| 38 | 30 \| 22 | 30 \| 33 |

As shown in **Fig. 2B**, an RF analysis compared metabolites between butyrate-treated WT and SOD1^G93A^ mice which lists the top 30 biochemicals that contribute to maximum importance. Metabolites classified as lipids such as oleoyl-linoleoyl-glycerol, lactosyl-N-palmitoyl-sphingosine, and lactosyl-N-stearoyl-spingosine had the highest degree of separation in the analysis. Additionally, other biochemical pathways showed decreasing separation, including (A) carbohydrate, (B) protein/amino acids, (C) lipid, and (D) xenobiotics metabolism.

#### Changes of carbohydrate in butyrate treated SOD1^G93A^ mice

As shown in **Fig. 2C**, SOD1^G93A^ mice showed a decrease in N-acetylneuraminate and N-glycolylneuraminate at 7 weeks of age following butyrate treatment (shown as But in figures). There is a decrease of N6-carboxymethyllysine (7 wk But vs Baseline, 14 wk vs Baseline), an advanced glycation end-product that can be utilized by colonic bacteria as a source of energy ^29^. These results suggest changes in carbohydrate metabolism following butyrate treatment. These differences appeared to be rectified in mice treated with butyrate. There were no time points of a statistically significant fold change ratio for the carbohydrate metabolites listed above except for pyruvate in which SOD1^G93A^ mice on average had a significant decrease compared to WT mice at 7 weeks of age. These results support the hypothesis that butyrate treatment helps rectify metabolomic alterations associated with ALS progression. Additionally, Ribulonate / xylulonate / lyxonate (isomers that co-elute together) were higher in ALS mice at baseline and decreased in mutant samples in response to butyrate administration (7wkBut vs Baseline, 14wkBut vs Baseline). Taken together, these decreases suggest that butyrate administrated to SOD1^G93A^ mice modified carbohydrate metabolism.

#### Changes of amino acids/protein in butyrate treated SOD1^G93A^ mice

In **Fig. 2D**, administration of butyrate resulted in lower levels of some tryptophan related metabolites in SOD1^G93A^ mice (7wk But vs. Baseline), which may be caused by enhanced uptake from the gut lumen and or increased degradation. ALS specific changes in tryptophan catabolites were observed in SOD1^G93A^ mice. The catabolites indoleacetate, indoleacrylate, kynurenate decreased at 7 weeks compared to baseline while at 14 weeks, anthranilate decreased. Other tryptophan catabolites including decreases in phenol sulfate, indolepropionylglycine, indoleacetylglycine and 3-indoxyl sulfate were observed (7 wk But vs Baseline, 14 wk But vs Baseline). The results suggest butyrate administration alters tryptophan catabolism in a genotype-dependent manner.

#### Changes of lipids in butyrate treated SOD1^G93A^ mice

Gut bacteria modulation of lipid metabolism is associated with weight gain in mice ^30^. **Fig. 2E** shows observed changes in sphingomyelins, sphingosines, and fatty acids. At 7 weeks post butyrate treatment, palmitoyl sphingomyelin, sphingomyelin, hydroxy palmitoyl sphingomyelin increased in SOD1^G93A^ mice while dehydrophyotosphingosine decreased at 14 weeks of age. These changes indicate fatty acids and phospholipid metabolism were altered in butyrate administration.

### Neuroactive metabolites increase significantly at onset in ALS mice

In the CNS, the neuroactive metabolites histamine and tryptophan play a crucial role in neuroprotection against neuroinflammation in SOD1^G93A^ mice ^31, 32^ and mediates inflammation in the PNS via production of mast cells in the gut ^33^ and producing commensal gut microorganisms via microbial decarboxylation of amino acids ^34^. Thus, we observed changes in histamine, tryptophan and glutamate related amino acids. In **Fig. 3A**, we showed a schematic of histamine metabolism and examined changes in histaminergic signaling at different time points using a SOD1^G93A^ / WT mouse model. The histidine catabolites 1-ribosyl-imidazoleacetate, 1-methyl-4-imidazoleacetate, 1- methylhistamine, histamine and N-acetyl-1-methylhistidine increased at 13 weeks of age in SOD1^G93A^ mice. In **Fig. 3B**, we illustrated a schematic of the tryptophan pathway and showed an increase in tryptophan catabolites N-acetyl tryptophan, N-formylanthranilic, 5-hydroxypicolinic acid, indoleacetate, indolepropionylglycine, indoleacetylglycine and 3- indoxyl sulfate for SOD1^G93A^ mice at 13 weeks of age. The last three metabolites had very significant increases at 13 weeks. At 17 weeks, N-acetyltryptophan and indoleacrylate decreased for SOD1^G93A^ mice. Taken together, these results suggest an alteration of neuroactive metabolites from the gut.

**Figure 3.**
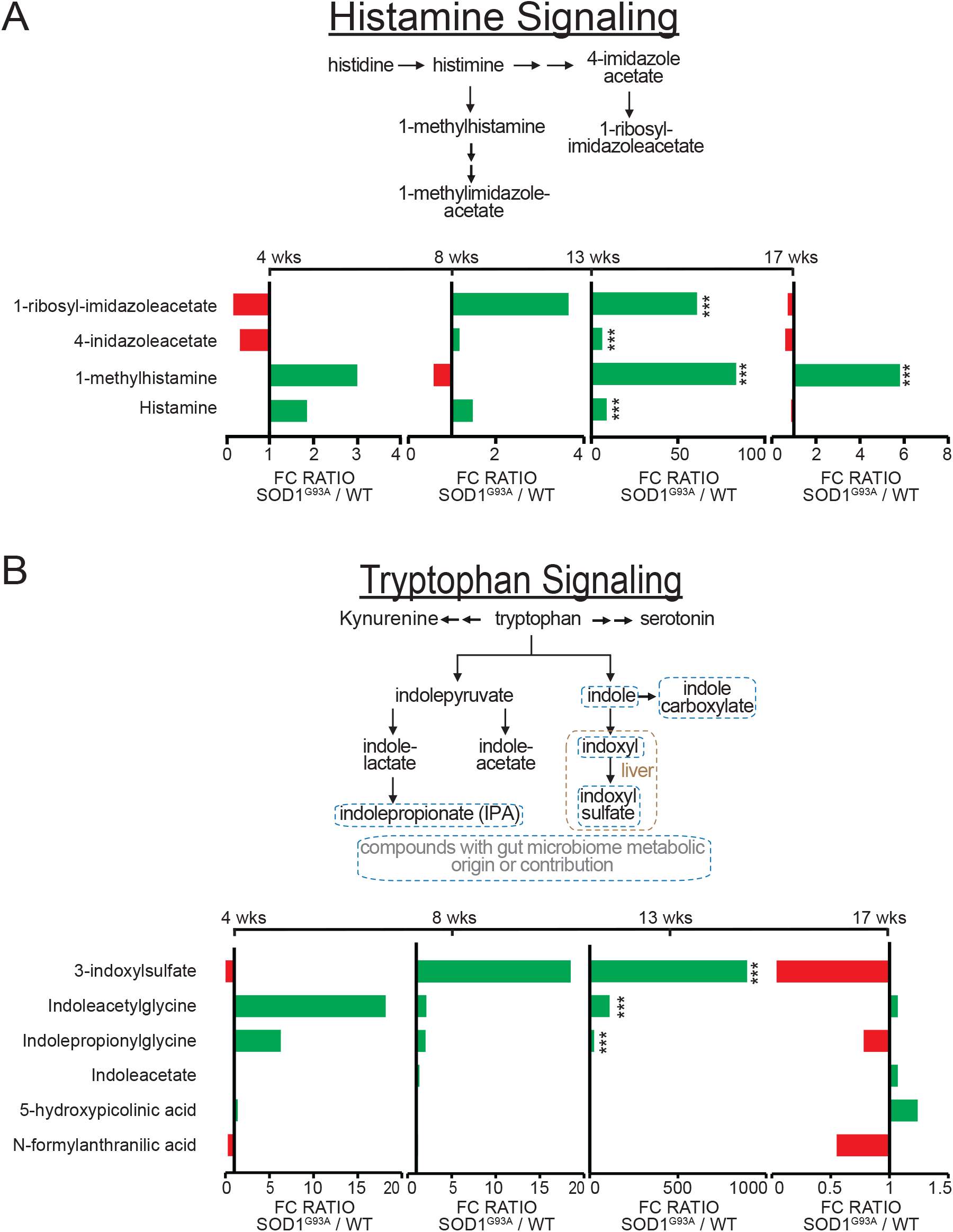
ALS altered histamine and tryptophan signaling at onset. (**A**) Schematic of histamine signaling, and longitudinal changes demonstrated by ALS mice over 17 weeks. (**B**) Schematic of tryptophan signaling, and longitudinal changes demonstrated by ALS mice over 17 weeks. Differences are assessed by the Welch’s two-sample t-test. WT (N= 6) & SOD1^G93A^ (N=6) mice. *P<0.05, **P<0.01, ***P<0.001.

### Dietary butyrate treatment altered neuroactive metabolites in ALS mice over disease course

We observed changes in histamine, tryptophan and glutamate metabolites following butyrate treatment in SOD1^G93A^ mice. An analysis of histamine catabolites showed a significant increase in histamine catabolites 1-methylhistamine and 4-imidazoleacetate at 7 weeks post-butyrate treatment (**Fig. 4A**). However, histamine catabolites 4- imidazole acetate and 1-methyl-5-imidazoleacetate decreased following butyrate treatment (baseline vs 7wk) (**Fig. 4B**). In **Fig. 4C**, we show tryptophan catabolite, indole- 3-carboxylate decreased after baseline in WT and SOD1^G93A^ mice (baseline vs. 7wk and 14 wk). Next, we observed changes in glutamate metabolism, in **Fig. 4D**, Glutamate, glutamine and N-methyl-GABA decreased following butyrate treatment while carboxyethyl-GABA only decreased for WT following butyrate administration.

**Figure 4.**
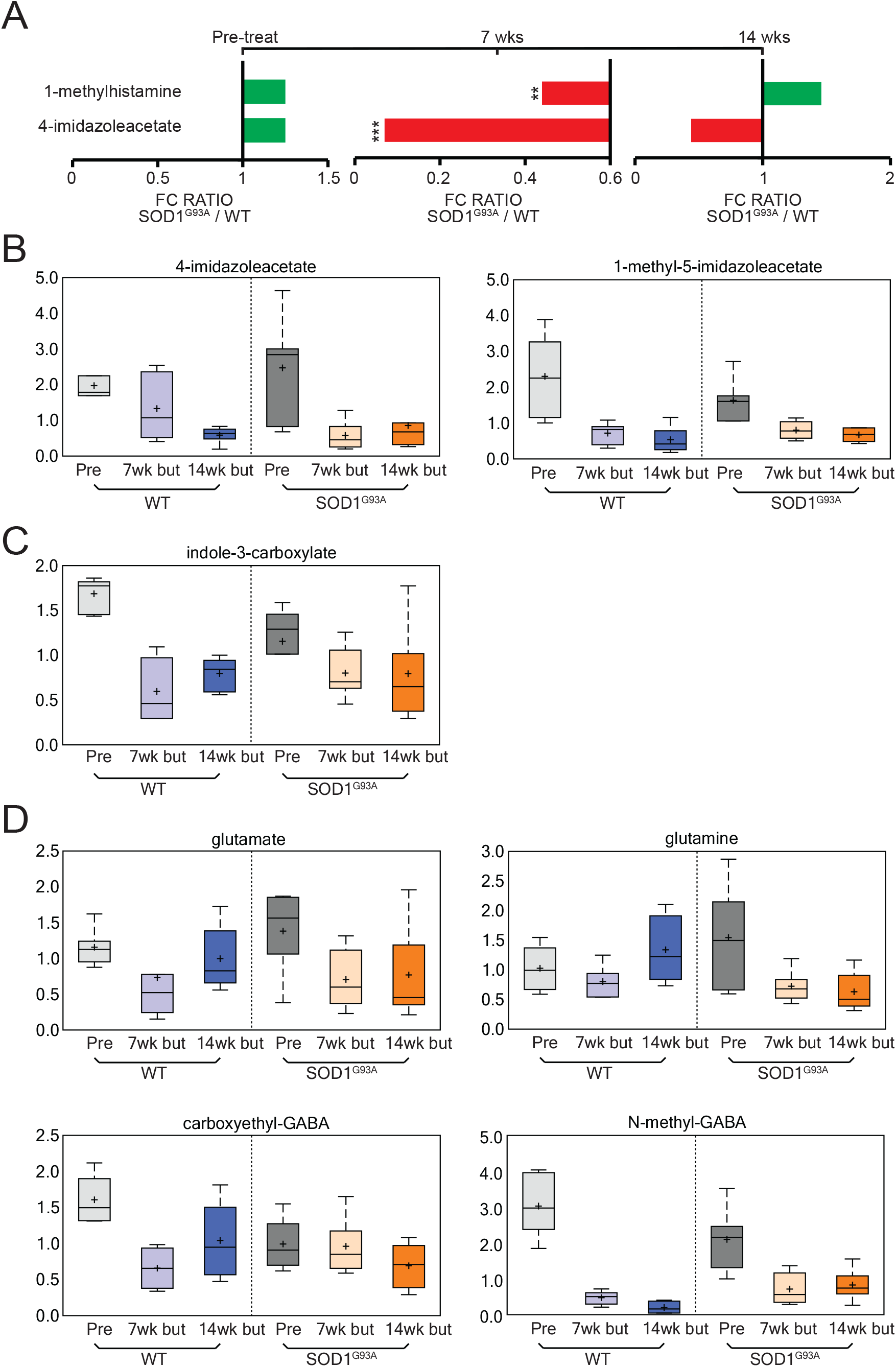
Butyrate treatment altered histamine and tryptophan signaling over the course of disease. (**A**) longitudinal changes of histamine signaling biochemical over the course of 14 weeks. Box-plot diagrams displaying longitudinal changes of butyrate-treated animals in (**B**) 4- imidazoleacetate and 1-methyl-5-imidazoleacetate. (**C**) indole-3-carboxylate (**D**) glutamate, glutamine, carboxyethyl-GABA, and N-methyl-GABA. The data presented as the fold change (FC) ratios of the average concentrations of WT and SOD1^G93A.^ Differences are assessed by the Welch’s two-sample t-test. WT (N= 6) & SOD1^G93A^ (N=6) mice. *P<0.05, **P<0.01, ***P<0.001.

### Energy related metabolites are altered during onset and decreased following butyrate treatment in ALS mice

Energy and mitochondrial dysfunction are involved in the pathogenesis of ALS. Animal and human studies have shown impaired mitochondrial metabolism that contributes to an increase in reactive oxygen species (ROS) ^35^. We assessed the changes in mitochondrial-associated metabolites during ALS. Citraconate/glutaconate, tricarballylate, alpha-ketoglutarate, aconitate, citrate and nicotinamide significantly increased at 13 weeks of age for untreated ALS mice (**Fig. 5A**). We investigated the effect of butyrate and found declines in nicotinamide and 1- methylnicotinamide levels were observed in response to butyrate treatment (14wk But vs Baseline, WT and SOD1^G93A^). Moreover, sarcosine, dimethylglycine and betaine were decreased in fecal samples collected from butyrate-treated mice (WT and SOD1^G93A^).

**Figure 5.**
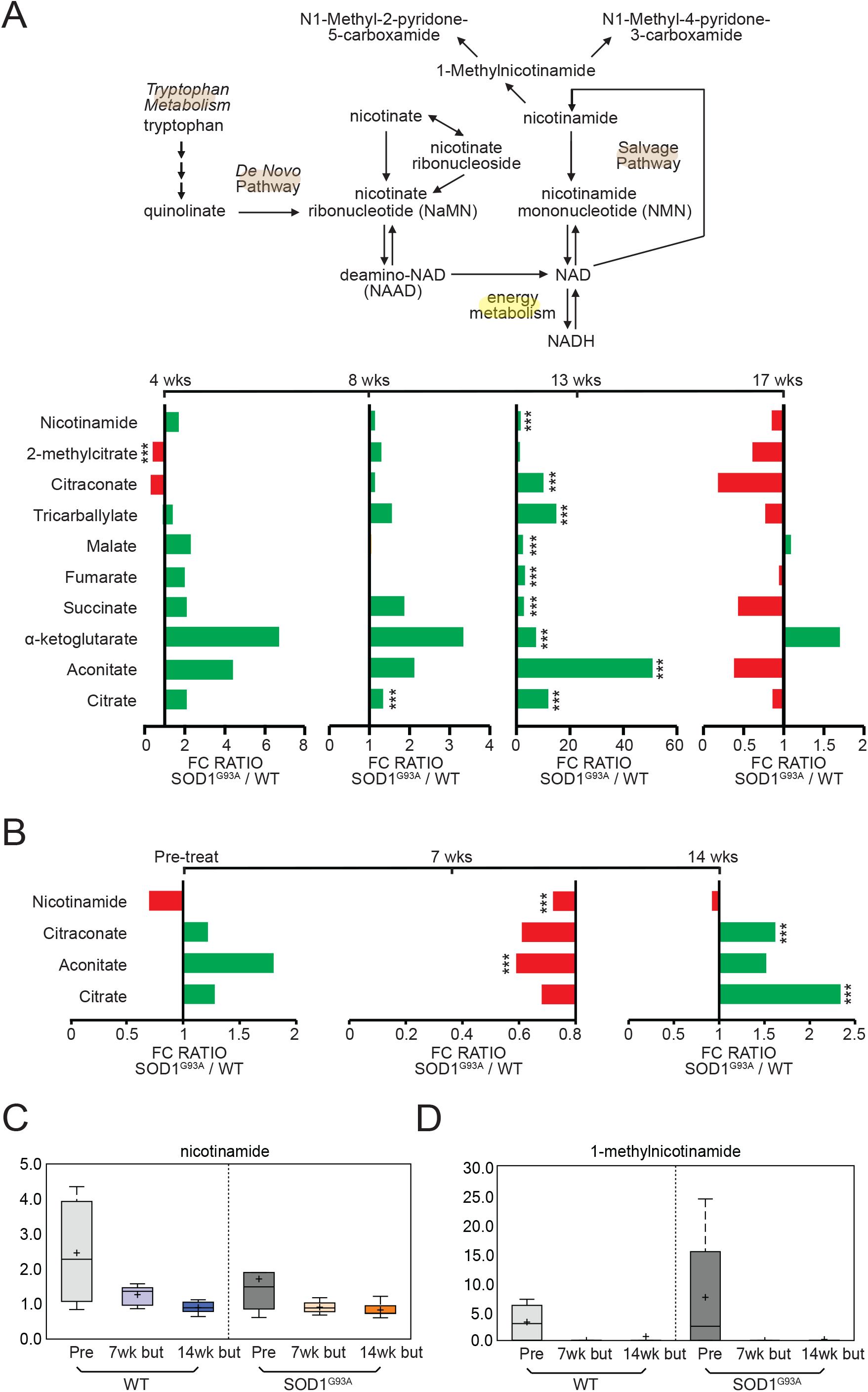
ALS altered energy-related metabolites over-time and butyrate treatment altered over the course of disease. (**A**) Longitudinal changes of energy-related biochemicals in ALS mice over 17 weeks. (**B**) Longitudinal changes of energy-related biochemicals in butyrate-treated ALS mice over 14 weeks. Energy-related metabolites (**C**) nicotinamide and (**D**) 1-methylnicotinamide have decreasing concentration. Differences are assessed by the Welch’s two-sample t-test. WT (N= 6) & SOD1^G93A^ (N=6) mice. *P<0.05, **P<0.01, ***P<0.001. * Significance is established at adjusted 0.05<P<0.1 and P-value<0.05.

These compounds are intermediates in one-carbon metabolism, which is implicated in nucleotide biosynthesis and methylation processes. Thus, these changes may potentially affect methylation pattern through modulation of substrate availability (**Fig. 5B**). In addition, nicotinamide and 1-methylnicotinamide had a decreasing trend for SOD1^G93A^ mice following butyrate treatment (**Fig. 5C and Fig. 5D**). These findings suggest butyrate treatment may influence energy metabolism.

### Gamma-glutamyl amino acid metabolites are altered following butyrate treatment in ALS mice

Gamma-glutamylated forms of amino acids are generated by gamma-glutamyltransferase (GGT), an enzyme that catalyzes the transfer of the glutamyl moiety from GSH to acceptor molecules (e.g. amino acids). Addition of gamma-glutamyl moiety improves the transport of amino acids across lipid membranes. Importantly, both host cells and certain bacterial species express GGT activity. In **Table 3**, we showed a schematic of the GGT pathway and detected declines in fecal levels of several gamma-glutamyl amino acids in butyrate-treated WT and SOD1^G93A^, which could be interpreted as a decrease in GGT activity. It is possible that changes observed here in response to butyrate administration may modify the neuronal function of SOD1^G93A^ mice.

**Table 3.** But treatment causes shifts in the gamma glutamyl amino acid profile in ALS, compared to WT mice.

| Biochemical Name | WT |  | G93A |  |
| --- | --- | --- | --- | --- |
|  | 7 wk but/<br>baseline | 14 wk but/<br>baseline | 7 wk but/<br>baseline | 14 wk but/<br>baseline |
| Gamma-glutamylalanine | 1.34 | 0.59 | 0.51 | 0.92 |
| Gamma-glutamylglutamate | 0.49 | 0.33 | 0.26 | 0.43 |
| Gamma-glutamylglutamine | 0.79 | 0.78 | 0.30 | 0.42 |
| Gamma-glutamylglycine | 0.86 | 0.49 | 0.39 | 0.69 |
| Gamma-glutamylhistidine | 0.48 | 0.50 | 0.33 | 0.41 |
| Gamma-glutamylisoleucine | 0.81 | 0.61 | 0.37 | 0.61 |
| Gamma-glutamylleucine | 1.30 | 0.91 | 0.36 | 0.64 |
| Gamma-glutamyl-<br>apha-lysine | 0.90 | 0.60 | 0.37 | 0.58 |
| Gamma-glutamyl-<br>epsilon-lysine | 1.39 | 1.84 | 0.37 | 0.46 |
| Gamma-glutamylmethionine | 1.43 | 1.16 | 0.25 | 0.60 |
| Gamma-glutamylphenylalanine | 0.88 | 0.62 | 0.30 | 0.57 |
| Gamma-glutamylthreonine | 0.98 | 0.71 | 0.35 | 0.69 |
| Gamma-glutamyltryosine | 0.73 | 0.72 | 0.33 | 0.75 |
| Gamma-glutamylvaline | 1.20 | 0.74 | 0.32 | 0.59 |
| Gamma-glutamylserine | 0.57 | 0.56 | 0.44 | 0.69 |

### Butyrate treatment reduced lumbar spine IBA1 expression, serum IL-17, and LPS expression in SOD1^G93A^ mice

Overexpression of microglia is categorized as neurotoxic and contributes to the upregulation of pro-inflammatory cytokines in ALS ^36^. We have shown manipulating the microbiome via butyrate treatment has decreased some inflammatory responses in SOD1^G93A^ mice ^17^. Thus, immunoflurescence analysis of microglia (IBA1) expression shows an increase in SOD1^G93A^ mice and significant decrease for butyrate treated SOD1^G93A^ mice (**Fig. 6A**). ALS is associated with an increase in inflammatory cytokines^9^. We observed serum IL-17 decreased in butyrate-treated SOD1^G93A^ mice compared to untreated SOD1^G93A^ and serum LPS increased in SOD1^G93A^ mice and decreased in butyrate-treated SOD1^G93A^ (**Fig. 6B and Fig. 6C**).

**Figure 6.**
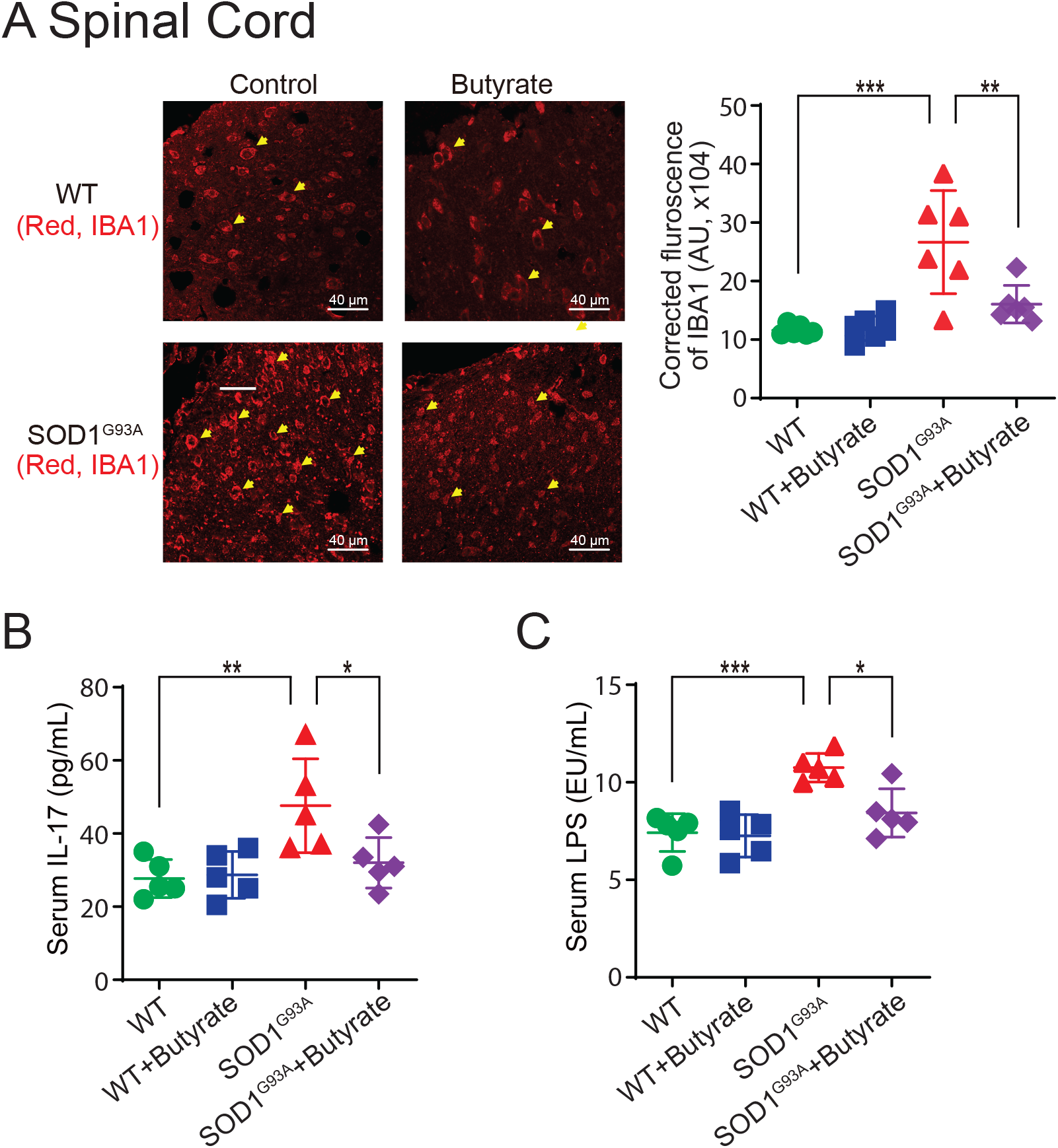
Butyrate treatment led to reduced lumbar spine IBA1 expression, serum IL-17 and LPS expression in SOD1^G93A^ mice. **(A)** Decreased IBA1 expression in lumbar spine of SOD1^G93A^ mice with butyrate treatment compared with the WT group. Images are from a single experiment and are representative of 6 mice per group. (Data are expressed as mean ± SD. n = 6, one-way ANOVA test, **P < 0.01, ***P < 0.001). **(B)** IL-17 was significantly lower in serum of SOD1^G93A^ mice with butyrate treatment compared with the WT group. (Data are expressed as mean ± SD. n = 5, one-way ANOVA test, *P < 0.05, **P < 0.01). **(C)** LPS was significantly lower in serum of SOD1^G93A^ mice with butyrate treatment compared with the WT group. (Data are expressed as mean ± SD. n = 5, one-way ANOVA test, *P < 0.05, ***P < 0.001).

## Discussion

In the current study, we investigated the longitudinal changes of fecal metabolites in the SOD1^G93A^ mice over the development of disease. We also compared the changes of metabolites in age-matched WT and SOD1^G93A^ mice treated with bacterial product butyrate. Samples were obtained from WT and SOD1^G93A^ mice at baseline and following 7 and 14 weeks of butyrate administration. In a parallel experiment, metabolic differences in feces from WT and SOD1^G93A^ mice were assessed in samples collected over the course of 13 weeks (4 weeks of age – 17 weeks of age). Individual samples were loaded in an equivalent manner across the analytical platforms and statistically analyzed. We identified changes in carbohydrate levels, amino acid metabolism, and formation of gamma-glutamyl amino acids. Understanding the metabolomics and intestinal changes will be important for the ALS pathogenesis and identification of biomarkers for diagnosis and targets for new treatments.

These compounds in the gut can be derived from digestion of plant hemicellulose by bacterial enzymes and the decreases observed here could suggest that butyrate administration resulted in altered composition/activity of intestinal microbiome. In fact, microbiota-induced changes in excretion of plant-derived carbohydrates have been reported before in mice subjected to feeding time restriction ^37^. This decrease suggest butyrate treatment could be mediating histamine levels through catabolites which could influence immune responses within the gut. Gut microbiota controls neurobehavior via modulating brain insulin sensitivity and metabolism of tryptophan, the precursor of serotonin ^38^. Bacterial cells can decarboxylate histidine to form histamine using histidine decarboxylase ^34^.

Carbohydrates play a crucial role in shaping the composition and physiology of the gut microbiome ^23^. **N-acetylneuraminate** and **erythronate** showed declines in response to butyrate treatment in samples collected from SOD1^G93A^ mice. These biochemicals can be produced by the gut microbiota from degradation of chitin and bacterial cell wall, which suggests that butyrate in the diet affected the gut microbiome community structure. Finally, both WT and SOD1^G93A^ groups exhibited decreases in **tricarballylate**, a microbial metabolite that can be derived from the TCA cycle intermediate aconitate (14wk But vs Baseline), and in **N6-carboxymethyllysine** (7wk But vs Baseline, 14wk But vs Baseline). Importantly, N6-carboxymethyllysine, an advanced glycation end-product, can be utilized by colonic bacteria as source of energy ^39^, which further indicates differences in microbiome activity over the course of the study. Together, these results are consistent with butyrate-induced alterations in composition of intestinal bacteria, which in turn modifies digestion of carbohydrates in the gut. Declines in fecal levels of several pentoses in response to butyrate may be interpreted as increased absorption of carbohydrates from the gut lumen, which consequently may contribute to prevention of weight loss in mutants.

ALS metabolite results suggest microbial metabolites could contribute to ALS symptoms at onset via glucose metabolism. Glucose metabolic dysregulation has been implicated in ALS ^24^. These fibers play a major role in shaping the gut microbiome composition and physiology. Past data indicated abnormalities in glucose metabolism within CNS of G93A mice.

Previous studies have highlighted imbalanced energy homeostasis in ALS patients ^40–42^ and transgenic ALS-expressing mice ^43, 44^. Certain metabolic alterations could contribute to the energy imbalance in ALS. Human studies have consistently found changes in glutamate metabolite levels by assessing the serum. One study examined the metabolite profile of serum from ALS patients via H NMR spectroscopy and found an increase in glutamate yet reduction of glutamine levels which indicates and imbalance in glutamate-glutamine conversion cycle ^45^. A different study used ^1^H NMR analysis to examine 17 metabolites in ALS patients and found a reduction of acetate and increase in acetone, pyruvate, and ascorbate concentrations ^46^. When comparing CSF metabolism between sALS and fALS patients, the results showed fALS patients had reduced glutamate and glutamine levels ^47^. The alteration of glutamate in ALS supports the hypothesis of glutamate toxicity. However, other metabolites are altered in ALS. An untargeted metabolomics study found post-diagnosed ALS patients have an increased benzoate metabolism, ceramide, creatine metabolism and fatty acid metabolism which suggest emerging pathways contribute to ALS ^48^.

Animal studies indicate alterations in the cerebrospinal fluid or the serum levels. A 2019 study examined the metabolites in the lumbar and thoracic spinal cord of two ALS mouse models and found alterations in lipid and energy related metabolites ^49^. Amino acid metabolites change during the course of ALS, Bame et al 2014 found amino acid profiles of ALS mice change between the ages of 50 and 70 days. Additionally, the amino acids: cysteine, methionine, glycine, sarcosine, phosphoserine and phosphoethanolamine were decreased in SOD1^G93A^ mice compared to WT. These metabolites are involved in the urea cycle, neuron excitation and methylation and related transsulfuration pathway ^50^.

Our data indicate the role of metabolites in neurophysiology. SOD1^G93A^ mice increased kynurenine, a pathway associated with inflammatory neurological disorder ^51^. SOD1^G93A^ mice displayed significant downregulation of quinolinate and tocopherol pathway derived metabolites and increase in nicotinamide. Quinolinic acid acts as a neurotoxin, proinflammatory mediator, and prooxidant molecule ^52^. The gut microbiota utilizes catabolism to ferment proteolytic amino acids as an energy source ^53^. The gut microbiota produces these metabolites via colonic fermentation of dietary fibers ^54^. Tryptophan is an essential amino acid that cannot be produced by the human body and must be obtained through diet. Defects in the tryptophan pathway is associated with development of diseases ^55^. The intestinal microbiome generates several metabolites from carbohydrates, amino acids, and other dietary sources. Several of these metabolites can bind to cellular receptors which could influence host functions ^34^.

Understanding metabolism in ALS through the gut microbiome is a growing field. We have shown the beneficial role of butyrate in treating the intestinal defects ^16^. A recent study has shown a direct impact of a gut microorganism on improving nicotinamide levels in ALS mice ^18^. Human studies also yield some similar findings. A human study used 50 ALS patients and 50 controls and found the gut microbiome changes during disease course yet ^56^. In contrast, a more recent study with 20 ALS participants found gene functions associated with metabolic pathways were reduced ^57^. Microbiota controls neurobehavior via modulating brain insulin sensitivity and metabolism of tryptophan, the precursor of serotonin ^38^. The role of the gut-neuron-microbiome axis needs further investigation in future research.

There are limited therapies to treat ALS patients. Our current study tried to explore the butyrate treatment for ALS by targeting metabolites. Paganoni *et al.* reported a phase 2 randomized, placebo-controlled trial involving 137 ALS patients, 89 of whom were treated with the combination of sodium phenylbutyrate plus taurursodiol ^58^. This combination has been shown to reduce neuron death and features of neurodegenerative diseases (including ALS) in preclinical models. This small trial, which treated patients for 24 weeks, found a modest reduction in functional decline in patients receiving the combination therapy ^58^. This is an encouraging finding, and larger trials testing more patients over a longer period are expected. The related changes of human microbiome and metabolites will help us to understand the efficacy and mechanism of the combination therapy.

The serum IL-17 and LPS were significantly reduced in the butyrate treated SOD1^G93A^ mice. Increased serum LPS is reported in Sporadic (SALS) patients ^8, 59^. SALS and familial (FALS) may have different initiating causes that lead to a mechanistically similar neurodegenerative pathway forms of ALS and clinical phenotypes. ALS patients have elevated intestinal inflammation and dysbiosis ^15, 60^. Some clinical studies have indicated intestinal abnormalities in ALS patients ^61–63^. Integrated “omic” biomarkers of microbiome and metabolites will help with the ALS diagnosis. Understanding inter-organ communications link among metabolites, inflammation, and ALS progression will provide novel insights into the potential to use metabolites as ALS hallmarks and for treatment.

## Conclusion

In conclusion, our study provides insights into fundamentals of metabolites in ALS. The results from this global metabolomic study assessing metabolic effects of butyrate administration on biochemical profiles of feces collected from WT and SOD1^G93A^ mice, together with longitudinally collected samples from untreated WT and mutant animals, differed in a number of metabolic readouts, including changes in carbohydrate levels, amino acid metabolism and formation of gamma-glutamyl amino acids. Differences in carbohydrates after butyrate treatment could be due to altered digestion and/or absorption of carbohydrates from the gut, which may be a consequence of changes in intestinal microbiota. Shifts is several microbially-contributed catabolites of aromatic amino acids are in agreement with butyrate-induced changes in composition of gut microbiome. Declines in gamma-glutamyl amino acids in feces may stem from differential expression of GGT in response to butyrate administration. Due to signaling nature of amino acid-derived metabolites, these changes could report on changes in inflammation (e.g. histamine) and contribute to differences in systemic levels of neurotransmitters (e.g. GABA, glutamate).

As a path forward, assaying plasma and brain samples from the butyrate experiment could allow characterization of systemic changes occurring after the treatment. Given that butyrate administration resulted in changes in NAD metabolism, which can affect the activity of sirtuins, and one-carbon metabolism, which can influence methylation processes, examining epigenetic marks in response to butyrate, may provide additional functional explanation for the protective effects of this compound in ALS. Novel therapeutic methods to target metabolites will improve the life quality of patients with ALS.

## Acknowledgements/Funds

We would like to acknowledge the VA Merit Award 1 I01BX004824-01, the NIDDK/National Institutes of Health grant R01 DK105118, R01DK114126, DOD CDMRP BC191198, and DOD BC160450P1 to Jun Sun. The study sponsors play no role in the study design, data collection, analysis, and interpretation of data. The contents do not represent the views of the United States Department of Veterans Affairs or the United States Government. Destiny Ogbu was a NIH PREP Scholar. Katerina Claud was a summer high school student and currently an undergraduate at Northwestern University.

## Author Contributions

DO and YZ, performed the animal studies, the detailed analyses of the results; KC and DO, perform the analyses of the metabolites and signaling pathways; DO and JS, prepared the figures and the draft text; YX contributed to the statistical analysis of data and the draft text; and JS obtained funds, designed the study, and directed the project. All authors contributed to the writing of the manuscript.

